# Quantifying the large-scale chromosome structural dynamics during the mitosis-to-G1 phase transition of cell cycle

**DOI:** 10.1101/2023.07.29.551121

**Authors:** Xiakun Chu, Jin Wang

## Abstract

Cell cycle, essential for various cellular processes, is known to be precisely regulated by the underlying gene network. Accumulating evidence has revealed that the chromosome, which serves as the scaffold for the gene expressions, undergoes significant structural reorganizations during mitosis. Understanding the mechanism of the cell cycle from the molecular chromosome structural perspective remains a grand challenge. In this study, we applied an integrated approach using a data-driven model combined with a nonequilibrium landscape-switching model to investigate large-scale chromosome structural dynamics during the mitosis-to-G1 phase transition. We generated 3D chromosome structural ensembles for the five critical stages in the process. We observed that the chromosome structural expansion and adaptation of the structural asphericity do not occur synchronously. We attributed this asynchronous adaptation behavior in the chromosome structural geometry to the unique unloading sequence of the two types of condensins. Furthermore, we observed that the coherent motions between the chromosomal loci are primarily enhanced within the topologically associating domains (TADs) as cells progress to the G1 phase, suggesting that TADs can be considered as both structural and dynamical units for organizing the 3D chromosome. Our analysis also reveals that the quantified pathways of chromosome structural reorganizations during the mitosis-to-G1 phase transition exhibit high stochasticity at the single-cell level and show non-linear behaviors in changing TADs and contacts formed at the long-range regions. These features underscore the complex nature of the cell-cycle processes. Our findings, which are consistent with the experiments in many aspects, offer valuable insights into the large-scale chromosome structural dynamics after mitosis and contribute to the molecular-level understanding of the cell-cycle process.

## 1 Introduction

Cell cycle, a fundamental and vital process, is crucial for cellular growth, proliferation, and development [1]. Cell cycle consists of four major, distinct phases: G1 phase, S phase, G2 phase, and the mitotic phase. The first three phases belong to the interphase, which occupies the majority of a cell’s time, while the last phase is related to the mitosis, involving the division of a parent cell into two identical daughter cells. Phase transitions in the cell cycle are precisely regulated by the underlying gene network, ensuring accurate genome replication and segregation to daughter cells during efficient cell cycle progression [2]. Currently, understanding the molecular mechanism governing the cell cycle process remains a grand challenge.

One of the most striking features of the cell cycle is the dramatic morphological changes in chromosomes as cells switch between interphase (decondensed structure) and the mitotic phase (condensed structure) [3]. Over the past two decades, rapid advancements in chromosome conformation capture techniques, particularly Hi-C methods [4, 5], have enabled the quantitative analysis of the chromosome structures at high spatial resolution. Hi-C measures the contact frequencies between the two DNA segments in the chromosome, resulting in a 2D contact map. Further examination of the Hi-C map can provide valuable insights into chromosome architecture. For example, Hi-C data from cells in the interphase often reveal insulated square blocks of elevated interaction frequency centered along the diagonal. This feature suggests the formation of topologically associating domains (TADs) [6, 7, 8, 9], where more frequent contacts are found within these megabase-sized domains than with neighboring regions. One functional advantage of TADs is to bring the distal enhancer and promoter sequences into the physical proximity, facilitating the gene expressions [10, 11]. At a higher level (> 5 Mb), Hi-C data display a plaid pattern, indicative of compartment formation [4, 5]. Chromosome compartmentalization is associated with the spatial segregation of gene-rich euchromatin and gene-poor heterochromatin. The compartment pattern, which varies by cell type, reflects structure-function regulation at long-range regions in chromosomes [12].

Accumulating Hi-C data on cell-cycle processes have revealed the disappearance of TADs and compartments during mitosis [13, 14, 15, 16, 17, 18]. Experimental data, along with theoretical models, consistently suggest cell-type-independent, cylinder-like chromosome structures during the mitotic phase [13, 15, 19], distinct from interphase chromosomes, which are considered fractal globules [20]. Consequently, large-scale chromosome structural reorganizations should occur during the cell cycle. Efforts have been made to perform time-course Hi-C during mitotic [15] and mitotic exit processes [16], as well as bulk Hi-C on highly purified, synchronous cell lines at different cell cycle phases [13, 17, 18]. Despite the wealth of Hi-C data, these experiments only offer a limited number of measurements on the temporal scale, failing to provide a continuous picture of cell-cycle chromosome dynamics. Moreover, Hi-C data only provide 2D information, lacking the 3D structure critical for understanding chromosome compaction and decompaction during the cell cycle. Thus, a reliable approach to investigate 3D chromosome structural dynamics at high spatial-temporal resolution is still needed.

We recently developed an integrated approach using a data-driven molecular dynamics (MD) simulation approach combined with a landscape-switching model, to study cell-cycle-dependent chromosome compaction and decompaction [21]. The model established a connection between the interphase and the mitotic phase, where Hi-C data are available, and generated continuous 3D chromosome structural evolution trajectories for the transitions between these two phases. Our approach, which is based on a coarse-grained model, overcomes the computational bottleneck for simulating the structural dynamics of the extremely long chromosome during the slow cellular processes. Furthermore, the theoretical predictions led to the consistent observations with the experiments in many aspects [13, 16], verifying the validity of the model. However, the simplified treatment of the cell cycle process only as a two-state transition between interphase and the mitotic phase has inevitably constrained the precision of the results [22]. Specifically, various pieces of evidence indicate that chromosome structures undergo dynamic reorganization throughout the cell cycle [17, 18, 15]. Therefore, conducting investigations while considering the chromosomal structural variations within the sub-phases of the cell cycle will be crucial in comprehending the accurate nuclear functionality during this fundamental cellular process.

In this study, we utilized Hi-C data from five critical stages during the transition from mitosis to the G1 phase [17], to refine the data-driven model and generate 3D chromosome structural ensembles at these five stages, correspondingly. We observed variations in the chromosome structural geometry at different stages from the prometaphase to G1 phase. These significant findings provide us with valuable opportunities to survey the molecular-level processes involved in organizing the 3D chromosomes during the mitosis-to-G1 transition. We then studied the transition between each pair of the adjacent cell stages using the landscape-switching model. We found that the coherent motions between chromosomal loci are enhanced in TADs throughout the transition, suggesting that TADs can be considered as both structural and dynamical units of chromosomes. We observed highly fluctuating chromosome structural reorganization trajectories for individual simulations, demonstrating that the chromosome dynamics are highly stochastic at the single-cell level [23, 14, 24, 25, 26]. Further analysis of quantified pathways revealed non-linear behavior in changing chromosome contact interactions during the mitosis-to-G1 transition. Notably, our results align well with the existing experimental evidence [15, 27], consolidating the validity of our findings. Overall, our study offers significant insights into the dynamics of large-scale chromosome structures after mitosis and enhance our understanding of the cell-cycle process at the molecular level.

## 2 Results

### 2.1 Chromosome structure reorganizations during the mitosis-to-G1 phase transition

Hi-C experiments have previously been conducted to investigate chromosomal structural reorganizations following mitosis [17]. These studies utilized highly purified, synchronous mouse erythroid cell populations at various stages: prometaphase (Prometa), anaphase or telophase (Ana/telo), early G1 phase (Early G1), mid G1 phase (Mid G1), and late G1 phase (Late G1). This allowed for the examination of chromosomal structures during critical stages in the transition from mitosis to G1 interphase (Figure 1A). At the Prometa stage, chromosomes exhibit a cell-type-independent Hi-C pattern, characterized by locally accumulated, non-specific contacts along the diagonal. This suggests a highly condensed structure, devoid of topologically associated domains (TADs) and compartments (Figure 1B). As cells transition into the G1 phase, short-range contacts become insulated at specific sites, while a plaid pattern gradually emerges at long-range regions, ultimately leading to the establishment of TADs and compartments.

**Figure 1:**
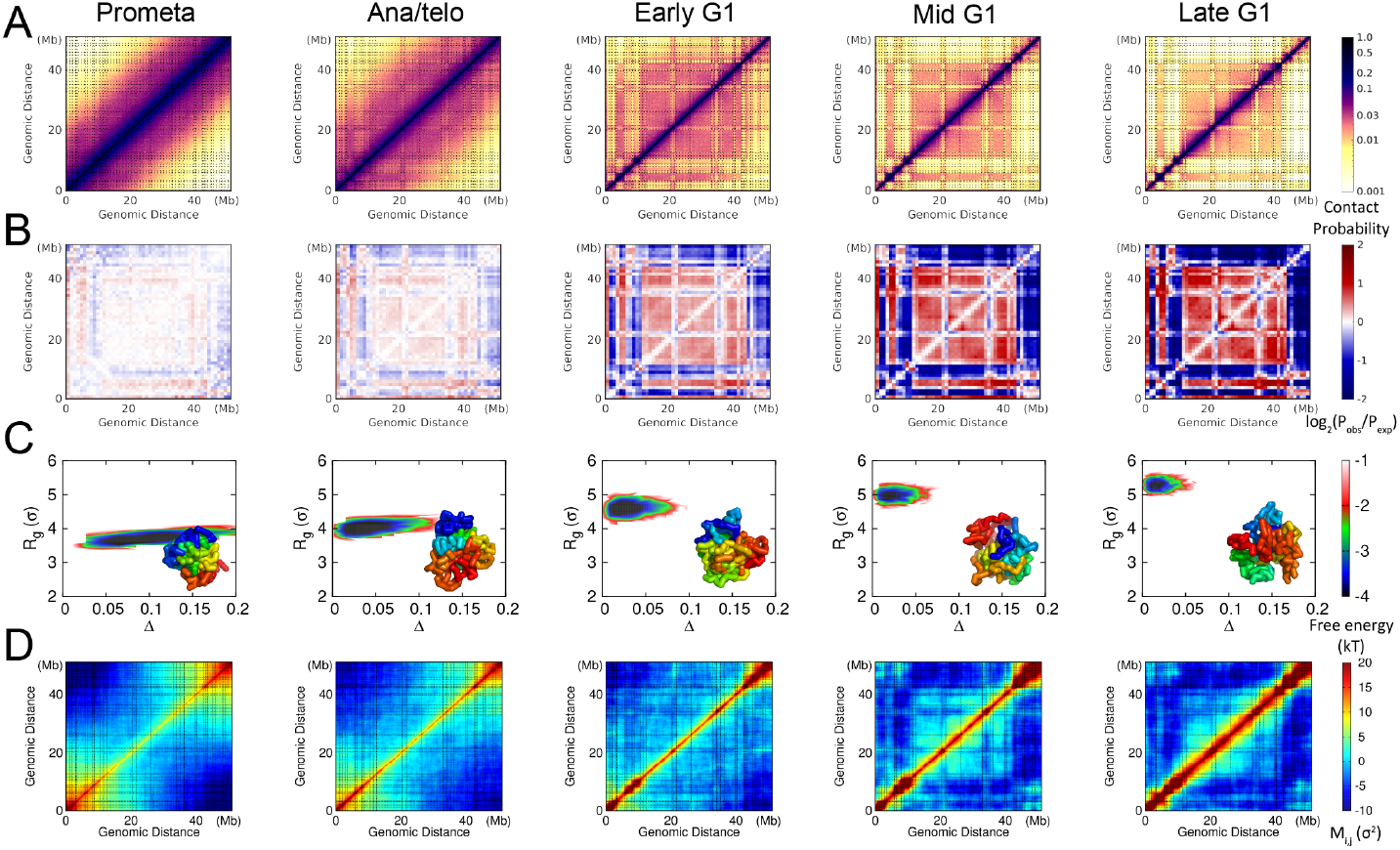
Chromosome structural properties at the Prometa, Ana/telo, Early G1, Mid G1 and Late G1 stages. (A) Hi-C contact maps. The contact matrics were normalized, so that the adjacent chromosomal loci (*i, i ±* 1) are always in contact (*P*_*i,i±*1_ = 1.0). (B) Enhanced contact maps. The enhanced contact probability was calculated as the ratio of the observed contact probability (*P*_*obs*_) over the expected contact probability (*P*_*exp*_) [4, 28]. (C) Free energy landscapes of the chromosome structural ensembles projected onto the geometrical quantities. The chromosome structural ensembles were generated by maximum entropy principle coarse-grained MD simulations. The geometry of the chromosome is described by the radius of gyration *R*_*g*_ and the aspheric shape quantity Δ. *σ* is the length unit of the coarse-grained model. Representative chromosome structure is shown in each panel. (D) Correlated fluctuating motions of chromosomal loci in the chromosome structural ensembles, measured by *M*_*i, j*_. The segment of chromosome 1 (20.5–71.4 Mb) was chosen in this study. The resolution of contact probability is 100 kb and each bead represents one 100 kb DNA segment in our coarse-grained model. The dashed lines in (A) and (D) indicate the TAD boundaries in the late G1 phase, identified by the method based on the insulation score profile [29].

We employed coarse-grained MD simulations combined with a data-driven approach to construct the chromosome structural ensembles at the Prometa, Ana/telo, Early G1, Mid G1, and Late G1 stages. We focused on a long segment of chromosome 1 (20.5–71.4 Mb) and modeled the chromosome with a bead-on-a-string representation at a resolution of 100 kb. The data-driven approach, which used the Hi-C data as restraints, was implemented through a maximum entropy principle strategy [30, 31, 19]. Subsequently, individual MD simulations were performed for each of the five stages, resulting in the generation of distinct chromosome structural ensembles.

Our coarse-grained MD simulations produced highly consistent contact maps with experiments for all these five stages (Figure S1-S5). The structures of the TADs and compartments, which are often characterized by insulation score profiles [29] and enhanced contact probabilities [4, 28], are also in excellent agreement with the experiments. These features indicate that our data-driven approach can faithfully recapitulate the experimental observations based on the 2D contact maps. Furthermore, the coarse-grained MD simulations generated the 3D chromosome structural ensembles, which can be used for examining the structural properties based on the Cartesian coordinates of the chromosomal loci.

We calculated the geometrical quantities of the chromosome structural ensembles at the Prometa, Ana/telo, Early G1, Mid G1, and Late G1 stages, individually (Figure 1C). Here, we used the radius of gyration *R*_*g*_ to describe the degree of chromosome structural compaction, and aspheric quantity Δ to describe the degree of chromosome structural asphericity. A perfect sphere has Δ = 0, and a deviation of Δ from 0 indicates the aspheric degree of the chromosome. We found that chromosomes at the Prometa stage have the highest degree of structural compaction with the lowest *R*_*g*_ among these five stages. Furthermore, the mitotic chromosomes exhibit highly aspheric geometry, reminiscent of a cylinder-like shape. These structural features of chromosomes at the Prometa stage are in line with the previous experimental observations [13, 15]. As cells undergo the transition from mitosis to G1 phase, chromosomes gradually expand their structure (increasing *R*_*g*_) and change to a sphere-like structural geometry (decreasing Δ towards 0). Interestingly, we found that although the chromosome structures are highly aspheric with a similar degree of compaction at the Prometa stage, the overall shape can be quite heterogeneous, demonstrated by a wide range of Δ. This further implies that the chromosome structural geometries are more diverse for different cells at the mitotic phase, compared to the ones at the interphase.

Chromosomes are highly flexible polymers. To investigate the structural flexibility of chromosomes at the five stages of cell transition from the mitosis to the G1 phase, we quantified the correlated fluctuating motions of chromosomal loci by calculating the fluctuation matrix (Figure 1D) [32, 33]:

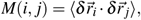

where 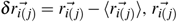 is the coordinate of chromosomal locus *i* (*j*), and 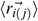 is the corresponding average. The fluctuation matrix *M*_*i, j*_ measures the correlation of spatial displacements between the chromosomal loci *i* and *j* with respect to their corresponding averages in the chromosome structural ensembles. The sign of *M*_*i, j*_ indicates whether the motion between the chromosomal loci *i* and *j* is positively or negatively correlated.

At the Prometa stage, chromosomal loci are positively correlated only when they are sequentially close, particularly within the two termini of the chromosome. Meanwhile, the magnitude of motion correlation between chromosomal loci decreases as their genomic distance increases, but it may increase subsequently when the evolving chromosomal loci are separated far away in sequence (e.g., chromosomal loci from two termini of the chromosome). Progressing to the G1 phase enhances the positive correlation of motions between chromosomal loci at short-range regions, and the pattern *M*_*i, j*_ at long-range regions roughly resembles that of the enhanced probability. At the Late G1 stage, *M*_*i, j*_ exhibits a large positive value when chromosomal loci *i* and *j* are within the same TAD, suggesting that the motions within TADs are strongly correlated in a positive manner. This indicates that TADs serve not only as the structural units in forming the 3D architecture of the chromosome, but also as the dynamic units in governing chromosome motions. Interestingly, we found that strengthening the enhanced contact probability can lead to a weakening of the correlation of motions between chromosomal loci at certain regions (e.g., 12.5–20.4 Mb and 36.3–39.8 Mb). This suggests that strong compartmentalization in chromosomes does not always result in strong correlated motions between chromosomal loci. Overall, we found that the coherent motions in chromosomes are weakest at the Prometa stage during the transition from mitosis to interphase. When cells undergo the transition to the Late G1 stage, the coherence of chromosome dynamics is enhanced both within TAD structures and at long-range regions.

### 2.2 Chromosome structural dynamics during the mitosis-to-G1 phase transition

Hi-C experiments provided contact-based 2D information on ensemble-averaged chromosome structure for cells at five discrete stages during the transition from the mitosis to the G1 phase. Hi-C-based data-driven MD simulations generated 3D chromosome structural ensembles and further formulated expressions for the effective energy landscapes, which govern the structures and dynamics of chromosomes within one cell stage during the transition. To bridge the gap between adjacent cell stages, we employed the landscape-switching model to simulate the chromosome structural dynamics during the cell mitotic exit processes [21, 34, 35, 36, 37, 38]. The landscape-switching model was applied between any two adjacent cell stages during the transition by instantaneously switching the effective energy landscape from one cell stage to another. This switching implementation provides the energy excitation necessary to drive the system for state transition, leading to nonequilibrium dynamics. The model, which is consistent with the nonequilibrium nature of cell-state transitions, has been found effective in reproducing many aspects of the experimental observations on various cell-fate-decision-making processes [21, 34, 35, 36, 37, 38].

The landscape-switching model generated chromosome structural dynamical trajectories, enabling investigations of continuous transition processes where only experimental Hi-C data at the initial and final states are available (Figure S6-S9). Additionally, one simulation represents the structural dynamics for one chromosome, resembling the transition trajectory at the single-cell level. To quantitatively describe the pathways, we projected the trajectories onto the geometrical quantities of chromosomes (*R*_*g*_ and Δ) (Figure 2). We found high stochasticity for the transition from the chromosome structural perspective, as no dominant pathways can be identified. Notably, these results are reminiscent of previous experimental observations, where deterministic paths connecting different cell states with significant fluctuations were characterized by single-cell Hi-C techniques for studying cell-cycle processes [14].

**Figure 2:**
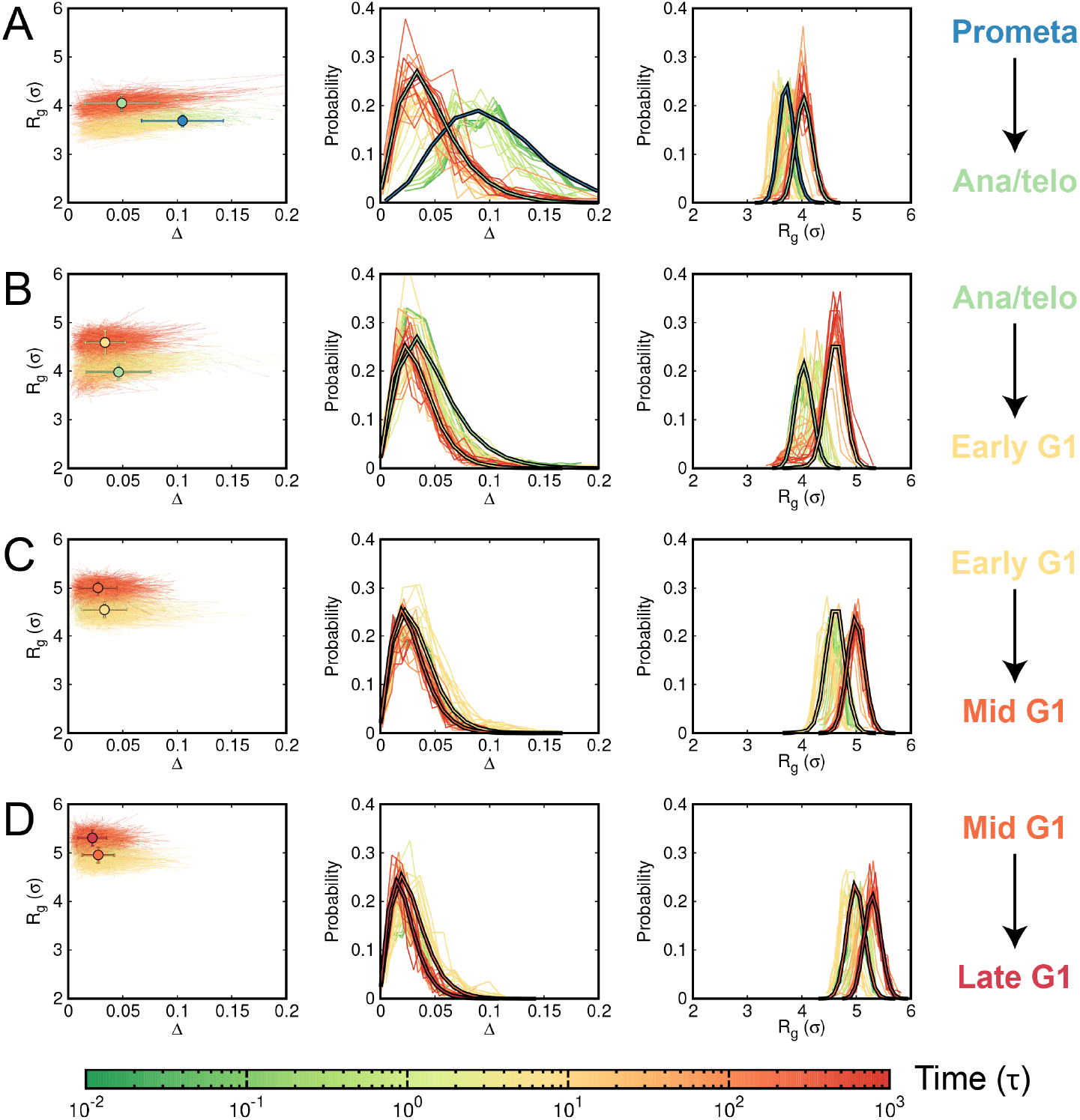
Geometrical properties of the chromosome structural ensembles during the transitions from mitosis to G1 phase, characterized by the radius of gyration (*R*_*g*_) and the aspheric shape quantity (Δ). (A) Transition from the Prometa to Ana/telo stage. (*Left*) Individual pathways with average values at the initial (Prometa) and final (Ana/telo) states shown as circles. (*Middle* and *Right*) Evolution of the probability distribution of Δ and *R*_*g*_ during the transition, represented by bold lines corresponding to the initial (Prometa) and final (Ana/telo) states. (B) Transition from the Ana/telo to Early G1 stage. (C) Transition from the Early G1 to Mid G1 stage. (D) Transition from the Mid G1 to Late G1 stage.

Considering the stochasticity of the trajectory, we calculated the evolution of the probability distribution of *R*_*g*_ and Δ during the transitions between every two adjacent stages. The trajectories clearly show that the geometrical quantities of chromo-somes adapt by changing the average values and distribution simultaneously during the transition. For instance, the most significant changes in Δ during the mitosis-to-G1 phase transition occur at the transition from the Prometa to Ana/telo stage, where the distribution of Δ gradually narrows down, accompanied by shifting the median towards smaller values. Although the asphericity of the chromosome structure largely adapts in the Prometa to Ana/telo transition, the expansion of the chromosome structure is not significant. The change in *R*_*g*_ reaches the highest degree in the transition from the Ana/telo to Early G1 stage, where only a minor change in Δ is observed. Adapting chromosome structural geometry in the progression of the G1 phase is mostly related to the further expansion of the chromosome structure, while the asphericity of the chromosome structure remains almost unchanged. Overall, we observed a significant adaptation of chromosome structural asphericity within mitosis from the Prometa to Ana/telo stage, while the expansion of the chromosome structure mostly occurs when the cell enters the G1 phase. The asphericity of the chromosome structure within the G1 phase is maintained while the structure continuously expands as cells proceed to the next stage of the G1 phase.

Our results indicate that the adaptation of the aspheric geometry and the structural expansion for chromosomes during the transition from the mitosis to the G1 phase are not synchronous. It has been well recognized that condensins play major roles in chromosome condensation during the mitotic phase [39, 40]. Numerous studies have revealed that two types of condensins (condensin I and condensin II), which form loops at different genomic distances [15], have distinct impacts on shaping the overall structures of chromosomes [41, 42]. Generally, condensin II promotes the formation of the long-range loops in the chromosome, while condensin I stabilizes and organizes the short-range loops, aiding in the lateral compaction of the chromosome [43, 15, 44]. Recent experiments showed that the absence of the condensin I during mitosis results in a disorganized, thicker chromosome scaffold, corresponding to low values of Δ. Conversely, without condensin II, chromosomes are unable to compact axially, leading to larger values of *R*_*g*_ [42]. Therefore, our findings indicate a sequence of condensin disassembly in which condensin I is unloaded first (decreasing Δ), followed by unloading of condensin II (increasing *R*_*g*_) during the transition from the mitosis to the G1 phase. The picture is consistent with the previous experimental observations, where in the telophase, condensin I is exported out of the reassembling nucleus whereas condensin II remains in the nucleus [45, 46].

To examine how the coherent motions between chromosomal loci evolve, we calculated the difference in the fluctuation matrix between the initial and final states during each transition (Figure 3). We observed that the changes in the coherence of the chromosome structural dynamics exhibit very different trends at different transitions. At the beginning of the transition from the Prometa to Ana/telo stage (10 *τ*), a noticeable decrease and increase in structural correlation within and between two termini of the chromosome were found, respectively. Meanwhile, *M*_*i, j*_ at the diagonal regions, which are related to TAD formations, remain largely unchanged. As the cell continues to proceed to the Ana/telo stage (after 10 *τ*), the structural correlations for the regions close in sequence enhance. In the transition from the Ana/telo to Early G1 stage, a non-specific, wide range of regions with increasing *M*_*i, j*_ was initially observed (10 *τ*), followed by a more specific pattern with increasing *M*_*i, j*_ at the local regions and between two termini of the chromosome at the late stage of mitosis (*t*=1000 *τ*). After the cell enters the G1 phase, the increase in *M*_*i, j*_ at the diagonal regions and between the two termini of the chromosome continues to be observed in the transition from the Early G1 to Mid G1 stage. Intriguingly, we observed that two distant regions belonging to the same compartment also exhibit an increase in *M*_*i, j*_, indicating a correlation of the chromosomal loci within the same compartment. The final transition to the late G1 stage leads to further enhancement of the coherent motions in the chromosome, primarily within the TADs. Therefore, we found that the correlation of chromosome structural dynamics within TADs is enhanced throughout the transition from the mitosis to the G1 phase. Since the boundaries of TADs have been found to be mostly established before entry to the G1 phase [17, 16, 22], the structural adaptations of TADs in the G1 phase are primarily related to strengthening the coherent motions within the domains, thus favoring their role as the basic unit for chromosome organizations.

**Figure 3:**
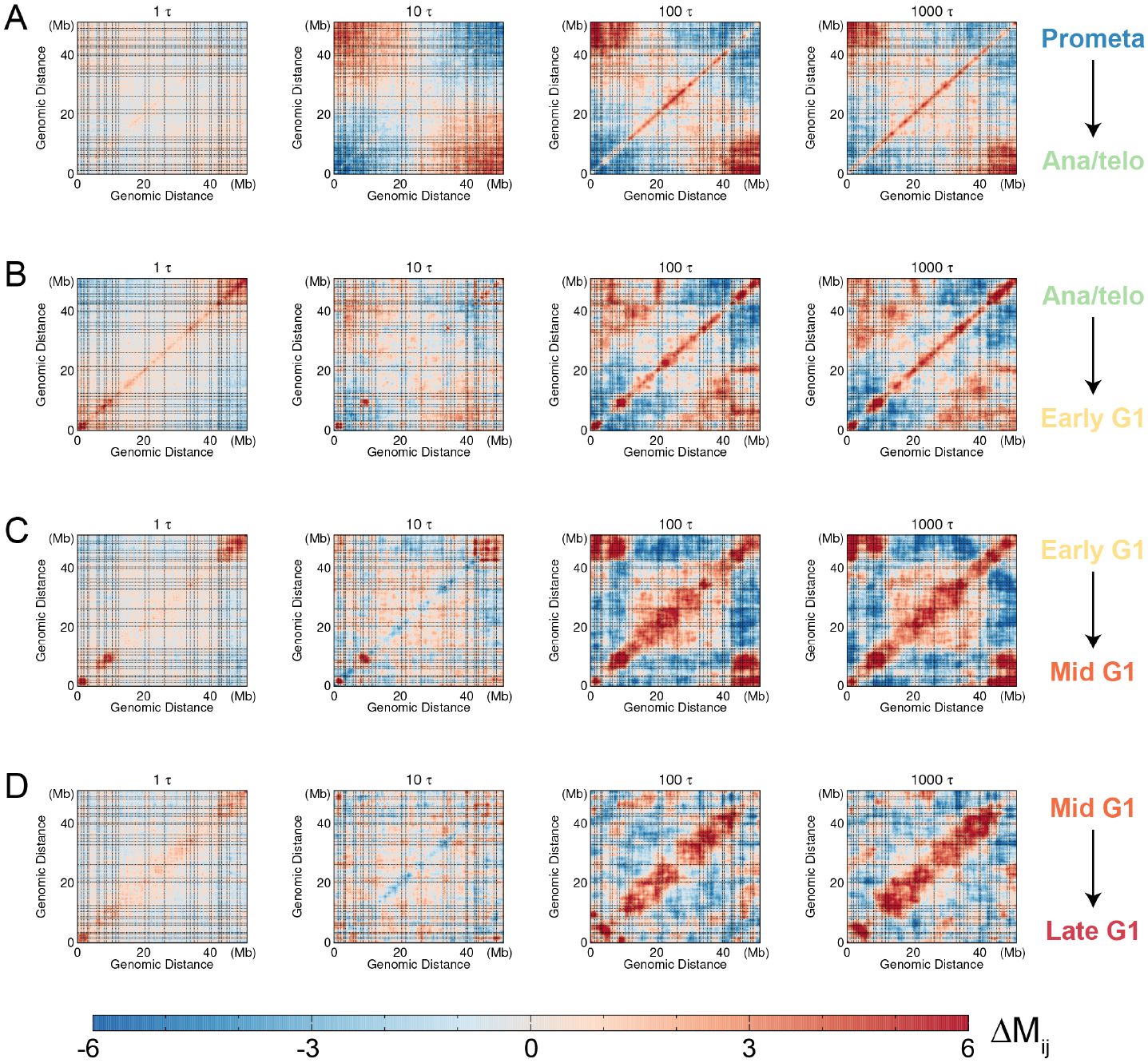
The difference in the fluctuation matrix between the chromosomal structural ensembles at the transition time *t* and the initial state for each transition. Δ*M*_*i, j*_ = *M*_*i, j*_ (*t*) − *M*_*i, j*_ (*t* = 0), where *T* represents 1*τ*, 10*τ*, 100*τ*, and 1000*τ*. (A) Δ*M*_*i, j*_ (*t*) for the transition from the Prometa to Ana/telo stage. (B) Δ*M*_*i, j*_ (*t*) for the transition from the Ana/telo to Early G1 stage. (C) Δ*M*_*i, j*_ (*t*) for the transition from the Early G1 to Mid G1 stage. (D) Δ*M*_*i, j*_ (*t*) for the transition from the Mid G1 to Late G1 stage. The dashed lines represent the TAD boundaries in the late G1 phase, as depicted in Figure 1.

### 2.3 Dynamics of TADs and compartments during the mitosis-to-G1 phase transition

As TADs have been identified as the structural units of chromosomes [47, 48, 49], we investigated the structural dynamics of TADs during the mitosis-to-G1 phase transition. We employed the insulation score profile, initially introduced to characterize TAD boundaries, to describe the structural formations of TADs. We conducted principal component analysis (PCA) on the evolution of the insulation score profile during each transition between the five stages during the mitosis-to-G1 phase transition (Figure 4A). We observed that the insulation score profiles of chromosomes at these five stages are distinctly separated, indicating structurally different TADs formed in these stages. This characteristic further suggests that TAD structures adapt throughout the transition from the mitosis to the G1 phase. Interestingly, the pathways observed deviate considerably from those obtained through linear interpolation of the Hi-C maps between the initial and final states of the transition. This implies that the landscape-switching model can produce non-linear behavior for the evolution of chromosome structure, potentially enabling the discovery of non-monotonic chromosome structural changes captured by previous experiments on cell-cycle processes [16].

**Figure 4:**
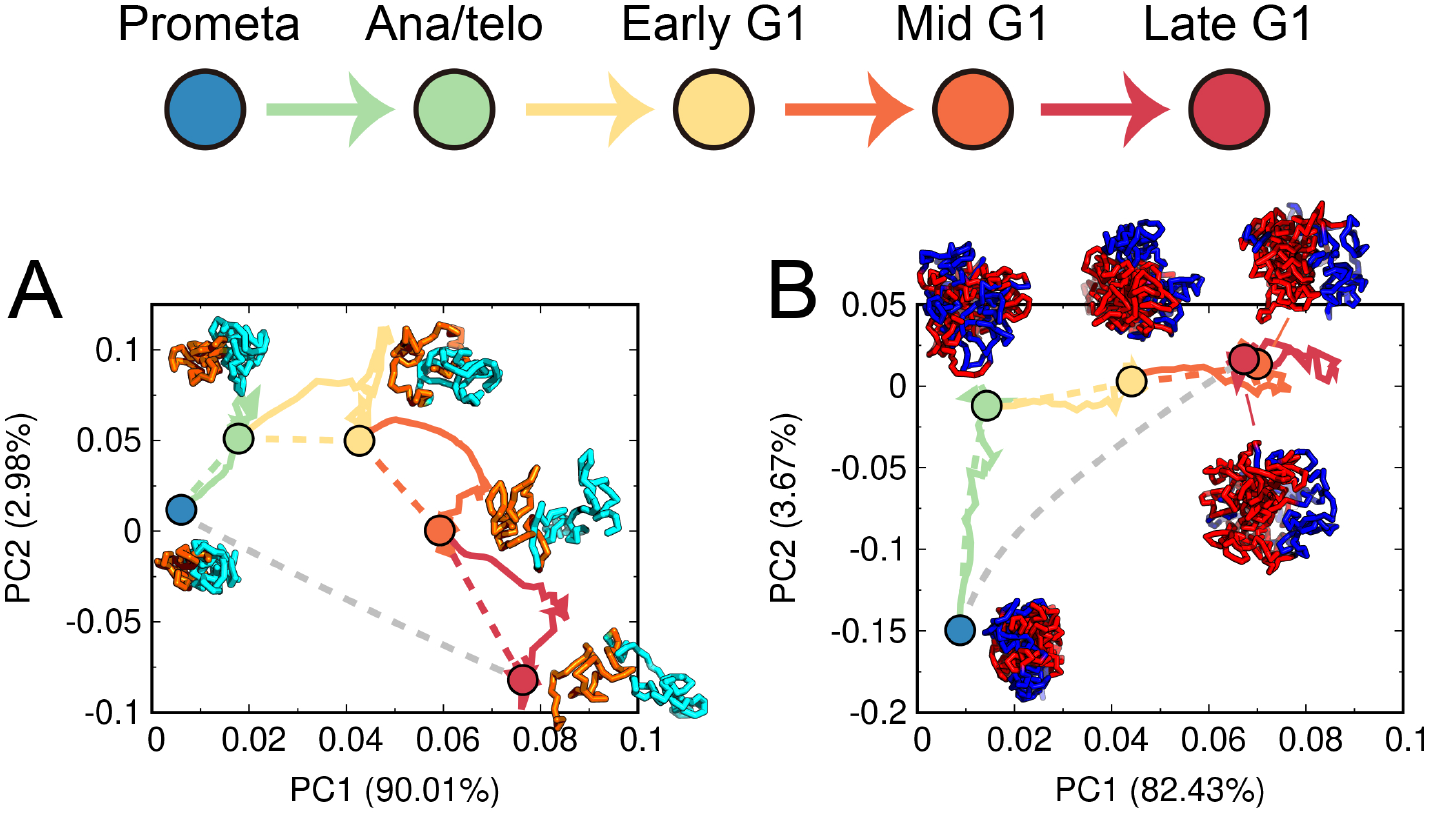
Quantified pathways of chromosome structural reorganizations during the transition from mitosis to G1 phase. (A) The quantified pathways (color solid lines) are shown in the PCA plot of the insulation score profiles projected onto the first two PCs. The color dashed lines indicate the pathways connected by the linear interpolation between the Hi-C data of the initial and final states for each transition. The grey dashed line indicate the the pathways connected by the linear interpolation between the Hi-C data of the Prometa and Late G1 stage. Representative structures of two consecutive TADs (21.6–26.2 Mb and 26.3–32.9 Mb), respectively colored by orange and cyan, are shown for these five cell stages. (B) The same as (A), but the PCA plot is done for enhanced contact probability *log*_2_(*P*_*obs*_*/P*_*exp*_). Representative structures of the chromosome with the locus colored by its corresponding compartmental annotation (red for compartment A and blue for compartment B), are shown for these five cell stages.

To examine the evolution of long-range structural organization in chromosomes during the transition from the mitosis to the G1 phase, we focused on the enhanced contact probability *P*_*obs*_*/P*_*exp*_, where *P*_*obs*_ and *P*_*exp*_ represent the observed and expected contact probability at 1 Mb resolution [4, 28]. Generally, the first principal component (PC1) of *P*_*obs*_*/P*_*exp*_ is used to describe the compartment profile of a chromosome, making *P*_*obs*_*/P*_*exp*_ suitable for describing long-range chromosome structural formation. We performed a similar PCA analysis on the evolution of *P*_*obs*_*/P*_*exp*_ during each transition (Figure 4B). We found that in the PCA plot, the Mid G1 and Late G1 stages are very close to each other and far from the other three distinct stages, suggesting that most long-range chromosome structural changes during the mitosis-to-G1 phase transition occur before the formation of the Mid G1 stage. Unlike TADs, we observed strong overlap between the pathways obtained from the landscape-switching model and the linear extrapolation method for each transition, indicating linear behavior for the evolution of chromosome structures at long-range regions during each transition. However, the linear connection between the Prometa and Late G1 stages deviates significantly from the pathways for the individual transitions within the process. These findings suggest that non-linear behavior of chromosome structural dynamics at long-range regions exists for the entire cell mitotic exit process, even though long-range contacts in chromosomes can be linearly interpolated by the initial and final states for individual transitions.

## 3 Discussion and conclusions

In this work, we investigated the chromosome structural dynamics during the mitosis-to-G1 phase transition using MD simulation approach with a combination of a data-driven model and the nonequilibrium landscape-switching model. We analyzed the chromosomal structural ensembles at five critical cell stages during the process, including prometaphase, anaphase or telophase, early G1 phase, mid G1 phase, and late G1 phase, as well as the transition pathways between adjacent cell stages. Our analysis was based on the geometric properties of chromosomal structures and the coherent motions between loci within chromosomes. We observed considerable fluctuations of the quantities in both the chromosomal structure ensemble and transition pathways, highlighting the stochastic nature of chromosomal structure and dynamics at the single-cell level [23, 14, 24, 25, 26]. Nevertheless, chromosomes undergo universal structural reorganizations, characterized by geometrical expansion, adaptation in aspheric shape, and enhanced coherent motions between chromosomal loci within the TADs during the transition from the mitosis to the G1 phase. These features suggest that chromosomal structural dynamics during the mitosis-to-G1 phase transition are driven by a combination of deterministic dynamics and stochastic effects, consistent with previous experimental findings on single-cell, cell-cycle-dependent Hi-C measurements [14].

Our results indicate that although chromosomes are highly condensed at the Prometa stage with the slowest diffusion dynamics of the chromosomal loci within a single cell among the five critical stages (Figure S1-S5), the degree of asphericity of the structure appears to span a wide range. This implies high geometrical variation for similarly compacted chromosomes (similar *R*_*g*_) in different cells at the Prometa stage. After mitosis, condensins dissociate from chromosomes, likely with simultaneous loading of cohesins [50], which are responsible for maintaining chromosome structure in the interphase [51]. However, the order of condensin-unloading and cohesion-loading during the transition from the mitosis to the G1 phase remains unclear. Our research, which sheds light on the complex dynamics of the chromosome structural geometry, may reveal insights into the molecular processes of the condensin unloading and cohesin loading after mitosis. A recent study used time-course Hi-C, chromatin binding assays and immunofluorescence experiments to identify an intermediate state for the condensin-to-cohesin transition after mitosis [15]. In this work, we further established that the unloading of condensin I precedes that of condensin II. This observation is based on the asynchronous changes of the chromosome structural geometry in terms of asphericity and expansion. Our findings have made a good contribution to the knowledge of the molecular-level mechanism in the transition from the mitosis to the G1 phase. By analyzing the coherent motions between chromosomal loci at the Prometa stage, we found highly positively and negatively correlated motions within and between the two termini of the chromosome, respectively. This suggests that the spatial motions of chromosomal loci at the Prometa stage are primarily related to structural compaction and decompaction, as these two termini move as individual wholes, in opposite directions. During the transition to the G1 phase, the compaction/decompaction motions at the chromosome termini gradually weaken, and the positively correlated motions between chromosomal loci within each TAD are progressively enhanced. In the G1 phase, the coherent motions between long-range regions are largely correlated with compartment formations. However, strong compartmentalization does not always result in a strong correlation in structural dynamics, leading to a decoupling relationship between the chromosome’s structure and dynamics. Based on this observation, we conclude that TADs, rather than compartments, can be considered as the fundamental units that encompass both the structural and dynamical aspects of chromosomes.

We quantified the chromosome structural reorganization pathways of TAD formations and the enhanced probability evolutions, which indicate the non-linear behavior of contact interaction reorganization in chromosomes during the mitosis-to-G1 phase transition. Notably, non-linear behaviors of chromosome structure dynamics related to the mitosis process have been widely observed in various experiments. For instance, Abramo et al. applied time-course Hi-C techniques to study the cell mitotic exit process and detected an intermediate state, where chromosomes are free of condensins and cohesins [16, 27]. They found that the chromosome contact interactions at this particular intermediate state differed significantly from the Hi-C data obtained using the interpolation-based method with the initial and final cell states as inputs, demonstrating non-linear behavior in modulating chromosome structures during the mitosis-to-G1 phase transition. This intermediate state was then successfully observed by our landscape-switching model, which further characterized the non-monotonic transition from the mitotic phase to interphase based on the quantified chromosome structural dynamical pathways [21]. Using live cell fluorescence imaging experiments, Chu et al. captured the shapes and sizes of chromosomes during mitosis at high resolution in 3D space and time [52, 53, 54]. They observed an interesting behavior that the sizes of chromosomes undergo an increase-followed-by-decrease non-monotonic adaption behavior as the cell proceeds from the prophase to cytokinesis. This is consistent with the results we obtained in our simulation. Although it seems to be popular, the functional role of this non-linear behavior in chromosome structural reorganizations after mitosis requires further investigations.

In summary, we investigated chromosomal structure ensembles of five critical cell stages during the mitosis-to-G1 phase transition. We quantified the pathways connecting each pair of adjacent cell stages during the process from the chromosomal structural perspective. Our model can be further improved by explicitly considering molecular-level processes [55]. Our landscape-switching model, which accounts for the nonequilibrium nature of cell-state transitions, predicted non-linear behavior of chromosomal structural reorganizations. This prediction provides valuable information for improving current interpolation-based methods widely used in 4D genome research [56, 57, 58].

## 4 Materials and Methods

### 4.1 Hi-C data processing

The Hi-C data for the murine erythroblastosis subline G1E-ER4 cells, at different cell cycle phases including prometaphase, anaphase, telophase, early G1 phase, mid G1 phase, and late G1 phase, were downloaded from the Gene Expression Omnibus database under accession number GSE129997 [17]. To analyze the Hi-C data, we utilized the Hi-C Pro software and followed the standard pipeline described by Servant et al. [59]. Furthermore, the Hi-C data was normalized using the iterative correction and eigenvector decomposition (ICE) method proposed by Imakaev et al. [60]. The resolution of the contact matrices was set to 100 kb. Our focus was on a specific region of chromosome 1, spanning from 20.5 Mb to 71.4 Mb, which contained a total of 510 beads in our coarse-grained beads-one-a-string model. To convert the Hi-C data into a contact probability map suitable for molecular dynamics (MD) simulations, we applied an additional normalization step by assuming that the highest contact probability was formed by neighboring beads with a value of *P*_*i, j*_ = 1.0, where *i* and *j* represent the chromosomal loci, and *j* = *i±* 1. This normalization approach enabled us to obtain a contact probability map for subsequent MD simulations.

### 4.2 Maximum entropy principle simulations

In our coarse-grained model, the structural and dynamical properties of the chromosome are governed by both bonded and non-bonded potentials. The neighboring chromosomal loci, represented by coarse-grained beads *i* and *i* ± 1, are connected through pseudo bonds, as described by Rosa et al. [61]. To account for the stiffness of the chromosome chain, we incorporated a linear-placement favored angle potential that acts on three adjacent beads. For the non-bonded interactions, we introduced soft-core repulsive forces between any pair of beads. This approach emulates the effects of topoisomerase enzymes, which play a role in untangling DNA chains [13, 31, 19]. Additionally, we included a spherical confinement to mimic the volume fraction of the chromosome within the cell nucleus at a 10%.

To incorporate the experimental Hi-C information into our coarse-grained chromosome model, we adopted a maximum entropy principle strategy, where the biasing potential should be in the linear form of the experimental observation [30]. Therefore, the overall potential *V* (*Stage*) can be expressed as the sum of two components: *V*_*Homopolymer*_ and *V*_*Hi*−*C*_(*Stage*). *V*_*Homopolymer*_ represents the non-specific homopolymer potential, which solely includes the bonded and soft-core non-bonded terms. On the other hand, *V*_*Hi*−*C*_(*Stage*) is the biasing potential that incorporates the experimental Hi-C data specific to the particular cell “Stage”. The expression for *V*_*Hi*−*C*_ is as follows:

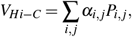

where *P*_*i, j*_ denotes the contact probability between the chromosomal loci “i” and “j,” and *α*_*i, j*_ acts as a prefactor governing the strength of the biasing potential. The values of *α*_*i, j*_ are determined iteratively through multiple rounds of MD simulation, aiming to minimize the discrepancy between the contact probabilities obtained from the simulated chromosome ensembles and the experimental data. For further details, please refer to the previous studies [31, 19, 62].

### 4.3 Landscape-switching model

We simulated the chromosome structural dynamics during the transition from mitosis to G1 phase using the landscape-switching model developed in our previous works [21, 34, 35, 36, 37, 38]. In brief, the model involved three main steps. First, we simulated the chromosome under the potential specific to the initial stage, denoted as *V* (*Stage*1). This potential was obtained through maximum entropy principle simulations incorporating the experimental Hi-C data. Then, we implemented a switching of the potential from the initial stage to the target stage, represented as *V* (*Stage*1) → *V* (*Stage*2). This switching process drives the system out-of-equilibrium. Finally, we simulated the chromosome under the new potential associated with the target stage, denoted as *V* (*Stage*2). The simulation allowed the system to relax on the post-switching energy landscape. The relaxation processes during this transition were collected and represented as the structural dynamical trajectories of the chromosome, capturing its conformational changes during the cell-state transition.

To reduce the large number of chromosome structures generated by the maximum entropy principle simulations, we performed structural clustering analysis on the ensembles. This analysis aimed to identify a smaller set of representative structures that could serve as initial structures for the subsequent landscape-switching MD simulations (Figure S1-S5). Within each cluster, we selected two chromosome structures, but only if the cluster’s population exceeded 0.2% of the ensemble. This selection process led to 170, 176, 176, 180, and 196 structures for the Prometa, Ana/telo, Early G1, Mid G1, and Late G1 stages, respectively. Subsequently, we generated the corresponding number of simulation trajectories for each stage transition, including the Prometa to Ana/telo stage, Ana/telo to Early G1 stage, Early G1 to Mid G1 stage, and Mid G1 stage to Late G1 stage. By employing this clustering and selection approach, we effectively decreased the number of chromosome structures at the specific cell stage, enabling us to focus on a reduced set of representative initial structures for the subsequent landscape-switching simulations.

### 4.4 MD simulation protocols

We utilized Gromacs (version 4.5.7) software [63] with PLUMED (version 2.5.0) [64], to conduct the MD simulations. The simulations were performed using Langevin dynamics with a friction coefficient of 10 *τ*^−1^, where *τ* represents the reduced time unit. In our simulations, the temperature was in the energy unit by multiplying the Boltzmann constant and *ε* is the reduced energy unit. Noteworthy, the temperature in the simulations does not directly correspond to real-world values. Instead, it represents the environmental scale that influences the structural dynamics of the chromosome under the specified potential [19]. The length was in the unit of *σ*, which represents the length of the pseudo bond in our model. Additionally, we used a time step of 0.0005 *τ* in the simulations to advance the dynamics of the system.

During each iteration of the maximum entropy principle simulations to calibrate *α*_*i, j*_, we conducted 100 independent MD simulations starting from different initial chromosome structures. Each simulation had a duration of 1000 *τ*. To enhance sampling of the conformational space, we employed a simulated annealing technique in these individual simulations. Initially, the temperature was gradually reduced from 4 *ε* to *ε* over the first 250 *τ* of the simulation. Subsequently, the temperature was held constant at *ε* for the remaining time. The second half of the trajectory, spanning from 500 *τ* to 1000 *τ*, was collected for the calculation of the simulated contact probability *P*_*i, j*_.

### 4.5 Structural and geometrical quantities

To assess the formation of TADs, we employed the insulation score introduced by Crane et al. [29]. Following the original study, we utilized a sliding window size of 500 × 500 kb to calculate the insulation score based on the contact map. The minima on the insulation score profile were then evaluated and identified as the boundaries of TADs [29]. To quantify the level of interaction enhancement, we calculated the enhanced contact probability, which was determined by dividing the observed contact probability *P*_*obs*_ by the expected contact probability *P*_*exp*_. The observed contact probability *P*_*obs*_ was obtained by summing the contact probabilities at a resolution of 100 kb within a 1 Mb region. On the other hand, the expected contact probability *P*_*exp*_ was calculated as the average contact probability between chromosomal loci separated by a specified genomic distance.

We used the radius of gyration *R*_*g*_ and aspheric quantity Δ to describe the geometry of the chromosome chain. These two quantities can be derived from the inertia tensor ***T*** with the following expression:

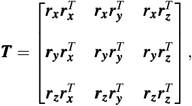

where ***r***_***x***_, ***r***_***y***_ and ***r***_***z***_ are the row vectors for the spatial positions of the chromosomal loci shifted by the corresponding means. *R*_*g*_ is expressed as follows:

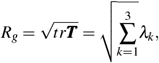

where Λ_*k*_ (*k*=1, 2 and 3) are the eigenvalues of ***T***, corresponding to the squares of the extension length along the three principal axes. Δ is expressed as follows:

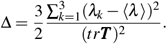

## Acknowledgement

We acknowledge the support from the National Science Foundation PHY-76066. The authors would like to thank Stony Brook Research Computing and Cyberinfrastructure, and the Institute for Advanced Computational Science at Stony Brook University for access to the high-performance SeaWulf computing system, which was made possible by a $1.4M National Science Foundation grant (#1531492).

## Notes

### Competing Interest Statement

The authors have declared no competing interest.

