## Supplementary material for "Quantifying the large-scale chromosome structural dynamics during the mitosis-to-G1 phase transition of cell cycle": SI Text

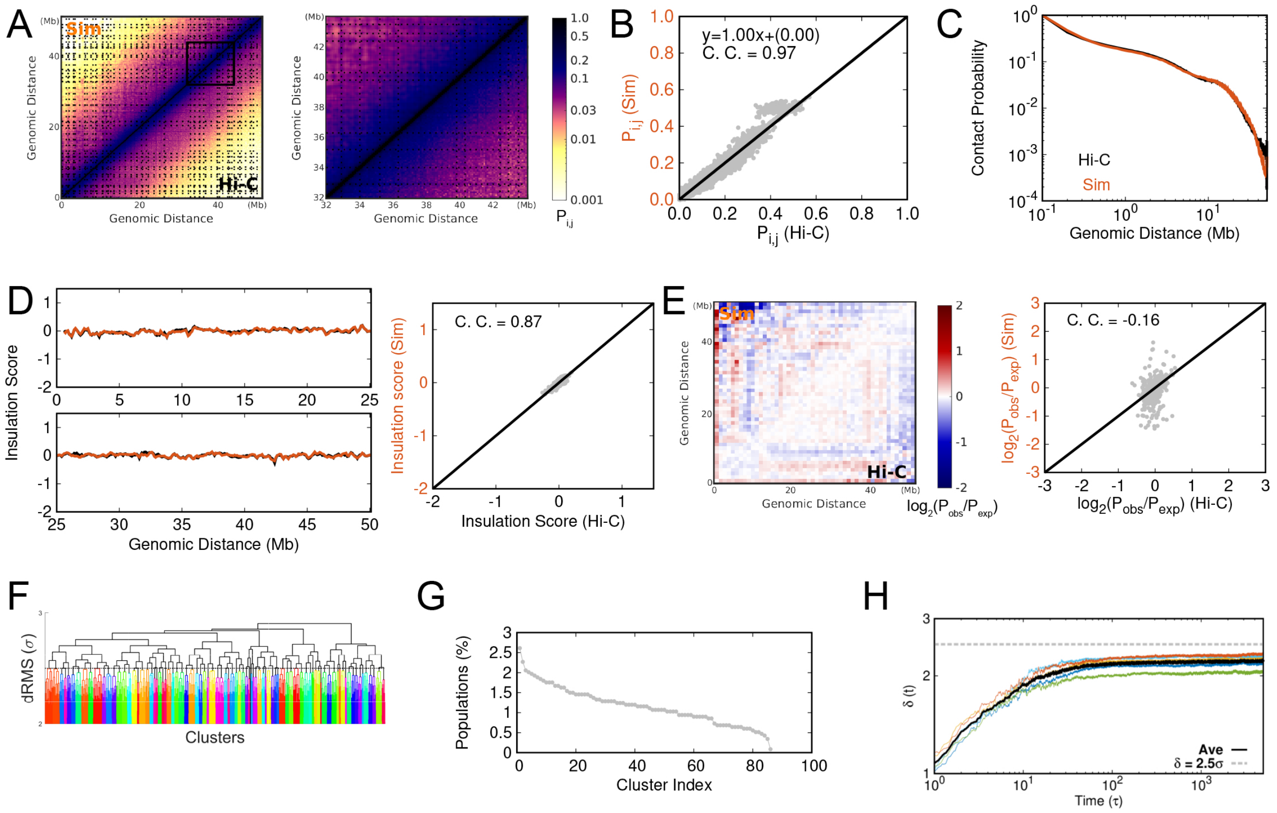


Figure S1: Chromosome structural ensemble and dynamics in the cell at the prometaphase (Prometa stage) and comparisons between the experimental Hi-C data and the data-driven MD simulations. The Hi-C data are publicly available at Gene Expression Omnibus (GEO) repository archives with accession number GSE129997 [1]. (A) Comparison of contact maps between the experimental Hi-C data and simulations. The *right panel* shows a zoomed-in contact map within the chromosomal distance range of 32–44 Mb. (B) Correlation of contact probabilities between the experimental Hi-C data and simulations. (C) Comparison of contact probability curves along the genomic distance between the experimental Hi-C data and simulations. (D) Comparison of insulation scores between the experimental Hi-C data and simulations [2]. (E) Correlation of enhanced contact probability $P_{obs}/P_{exp}$ between the experimental Hi-C data and simulations [3, 4]. (F) Dendrogram representing hierarchical clustering of the chromosome structural ensemble. (G) Population distribution of the clusters. (H) Time evolution of the average root-mean-square deviation of pairwise distances $d_{rms}$ between chromosomal locus pairs at time $t$ relative to their initial value $\delta(t) = \sum_{i,j} d_{rms}(i,j,t)/N_{pairs}$, where $N_{pairs}$ is the number of summed chromosomal pairs $i$ and $j$. Five trajectories are shown with different colors, and the black line represents the average. The dashed line indicates the threshold ($2.5\sigma$) for $\delta(t)$, which is used for structural clustering in (F), resulting in 85 clusters.

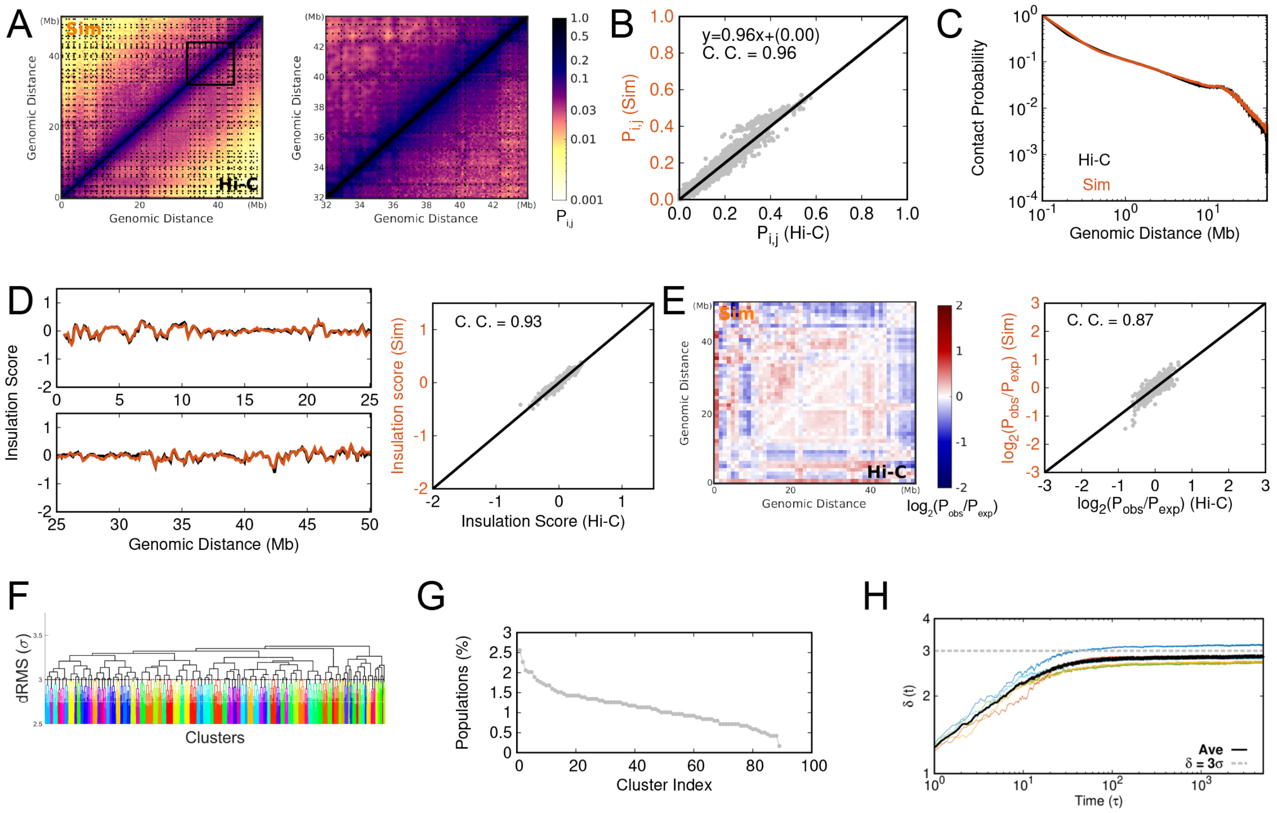


Figure S2: Same with Figure S1 but for the cell at the anaphase or telophase (Ana/telo stage). The dashed line in (H) indicates the threshold ($3\sigma$) for $\delta(t)$, which is used for structural clustering in (F), leading to 88 clusters.

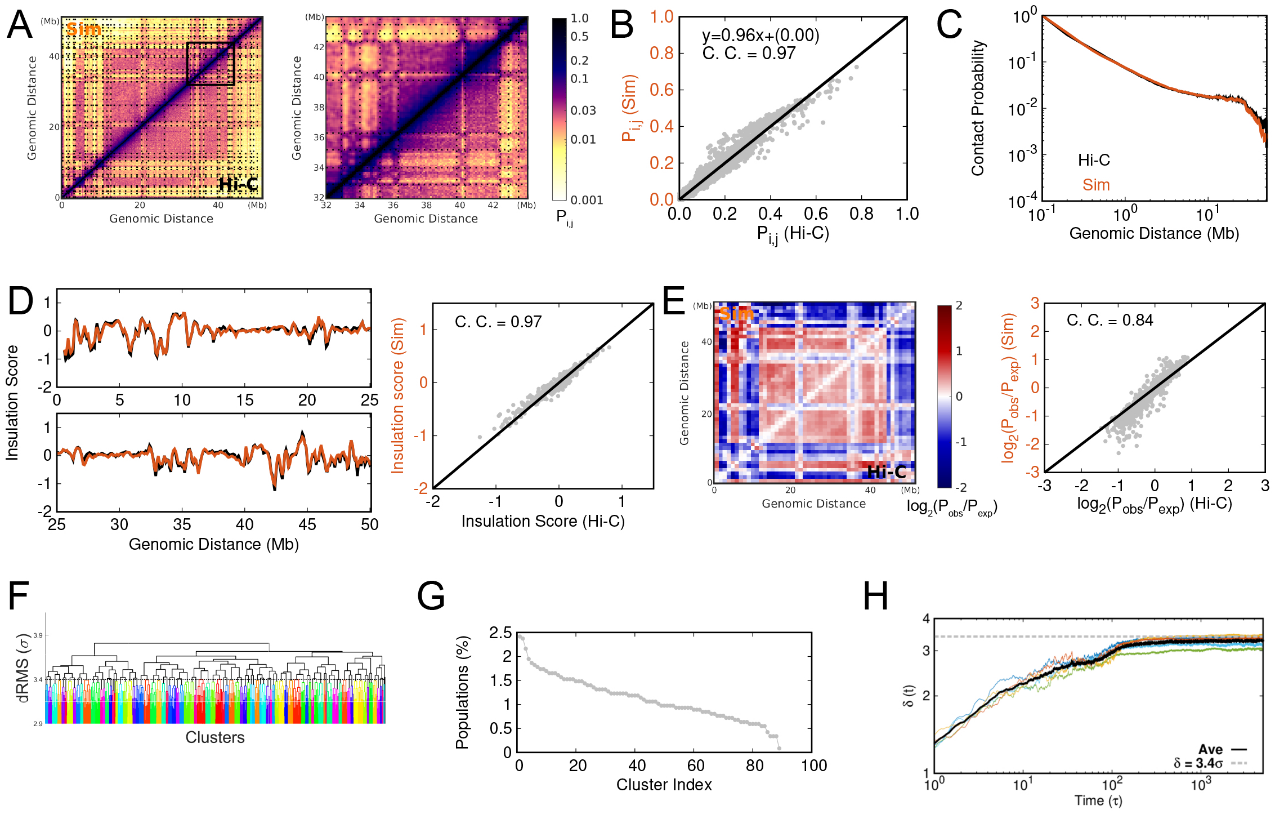


Figure S3: Same with Figure S1 but for the cell at the early G1 phase (early G1 stage). The dashed line in (H) indicates the threshold ($3.4\sigma$) for $\delta(t)$, which is used for structural clustering in (F), leading to 88 clusters.

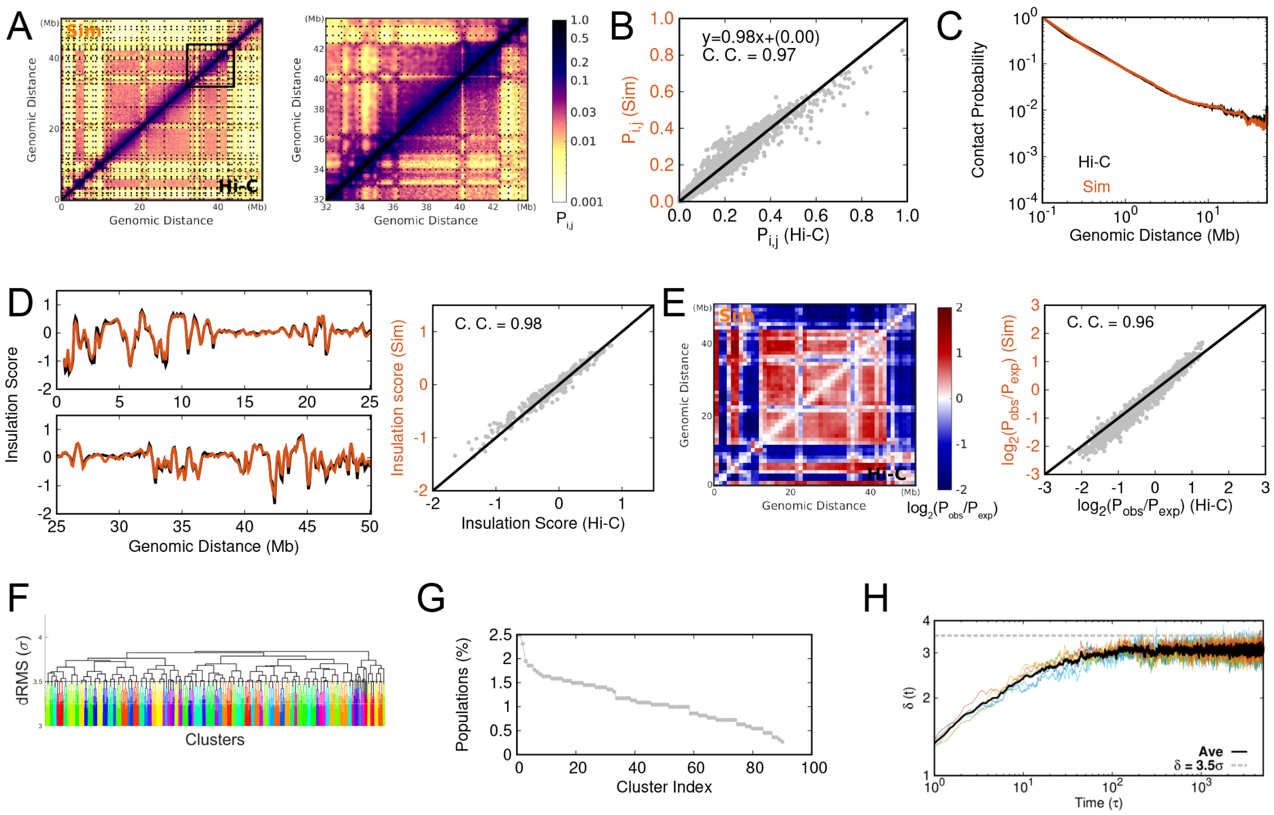

Figure S4: Same with Figure S1 but for the cell at the mid G1 phase (mid G1 stage). The dashed line in (H) indicates the threshold ($3.5\sigma$) for $\delta(t)$, which is used for structural clustering in (F), leading to 90 clusters.

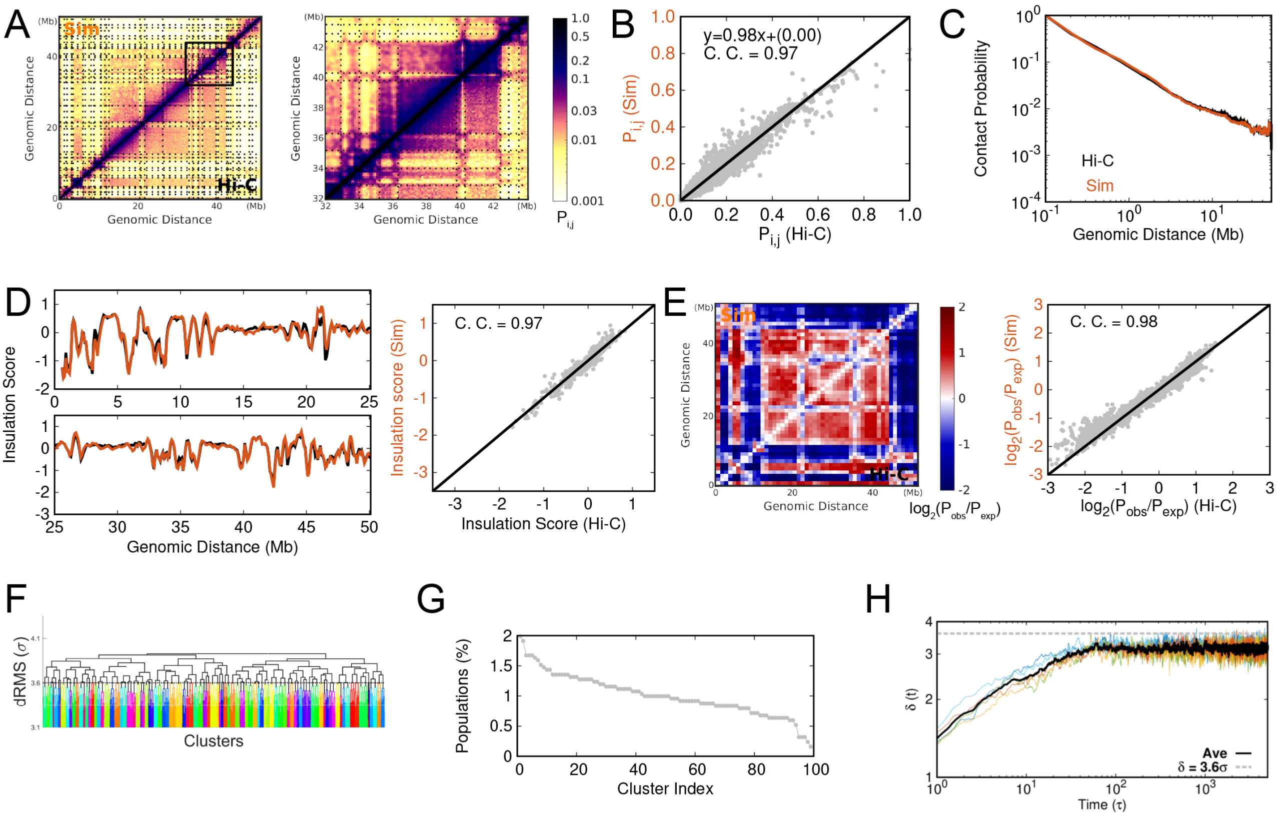


Figure S5: Same with Figure S1 but for the cell at the late G1 phase (late G1 stage). The dashed line in (H) indicates the threshold ($3.5\sigma$) for $\delta(t)$, which is used for structural clustering in (F), leading to 98 clusters.

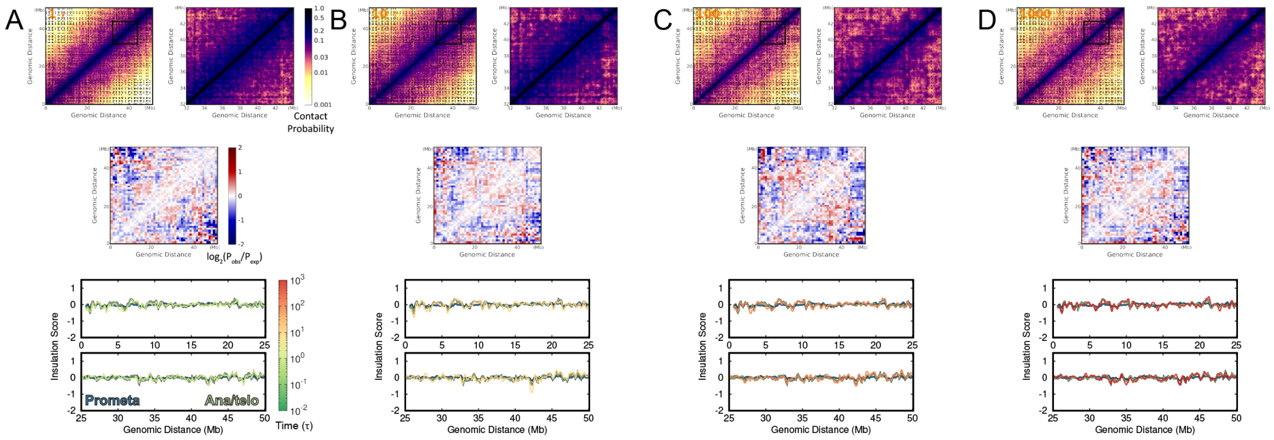


Figure S6: Structural evolution of chromosomes during the transition from the Prometa to Ana/telo stage. The contact probability maps, enhanced contact probability map, insulation score profile are shown at (A) 1 $\tau$, (B) 10 $\tau$, (C) 100 $\tau$ and (D) 1000 $\tau$.

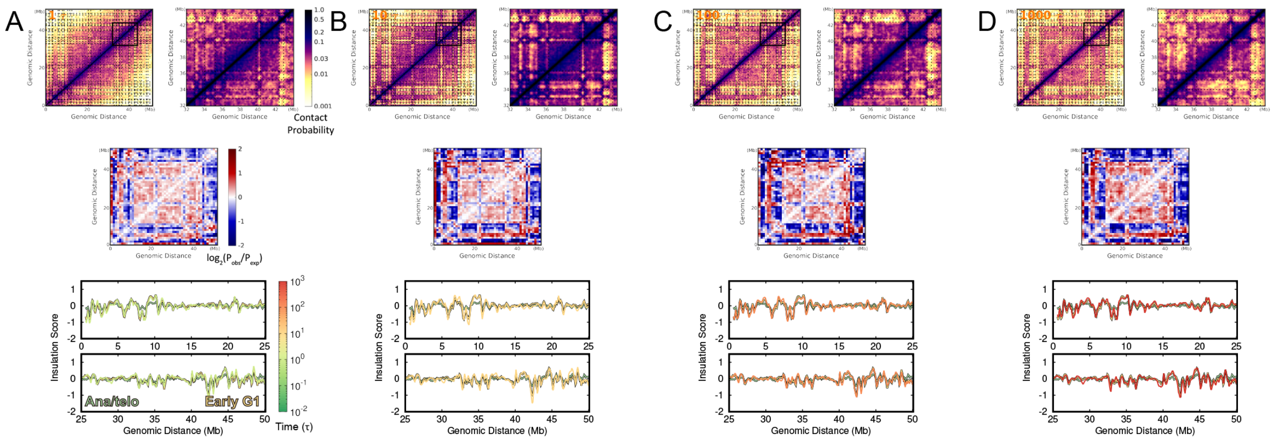


Figure S7: Structural evolution of chromosomes during the transition from the Ana/telo to Early G1 stage. The contact probability maps, enhanced contact probability map, insulation score profile are shown at (A) 1 $\tau$, (B) 10 $\tau$, (C) 100 $\tau$ and (D) 1000 $\tau$.

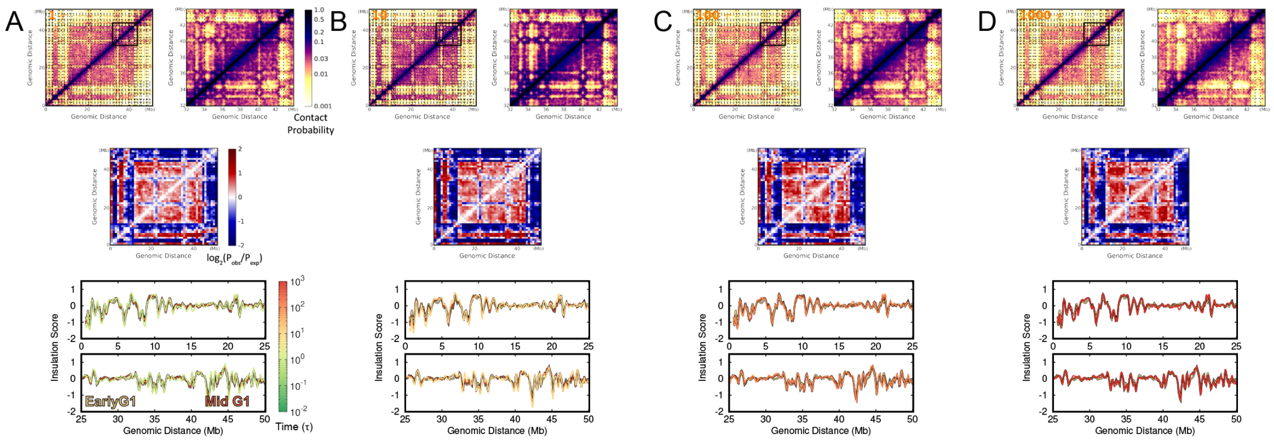


Figure S8: Structural evolution of chromosomes during the transition from the Early G1 to Mid G1 stage. The contact probability maps, enhanced contact probability map, insulation score profile are shown at (A) 1 $\tau$, (B) 10 $\tau$, (C) 100 $\tau$ and (D) 1000 $\tau$.

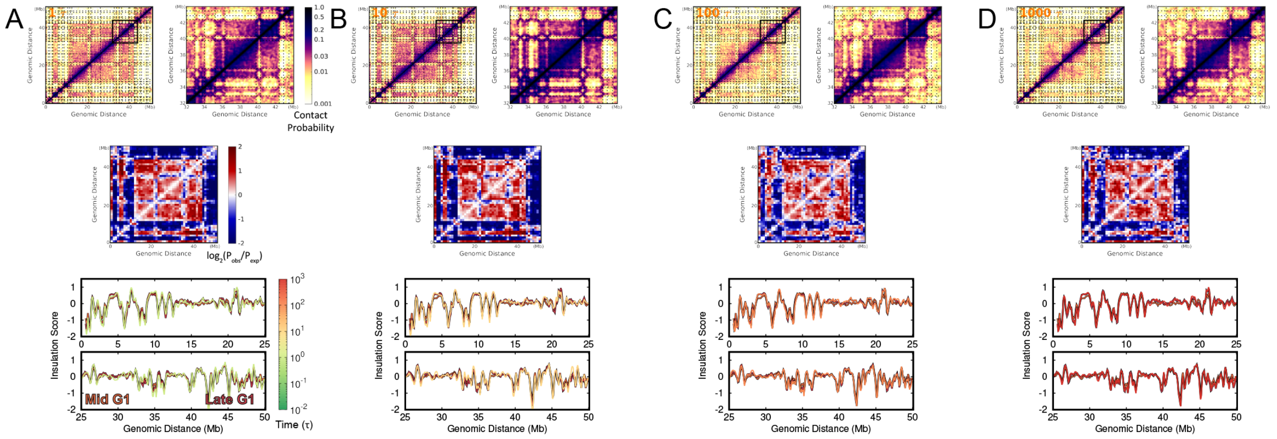


Figure S9: Structural evolution of chromosomes during the transition from the Mid G1 to Late G1 stage. The contact probability maps, enhanced contact probability map, insulation score profile are shown at (A) 1 $\tau$, (B) 10 $\tau$, (C) 100 $\tau$ and (D) 1000 $\tau$.
